# *Tm-2^2^* resistance targets a conserved cysteine essential for tobacco mosaic virus (TMV) movement

**DOI:** 10.1101/2022.10.06.510505

**Authors:** Hagit Hak, Hagai Raanan, Shahar Schwarz, Yifat Sherman, Savithramma P Dinesh-Kumar, Ziv Spiegelman

## Abstract

The tomato *Tm-2^2^* gene was considered one of the most durable resistance genes in agriculture, protecting against viruses of the *Tobamovirus* genus, such as Tomato mosaic virus (ToMV) and Tobacco mosaic virus (TMV). However, an emerging tobamovirus, Tomato brown rugose fruit virus (ToBRFV), has overcome *Tm-2^2^*, damaging tomato production worldwide. *Tm-2^2^* encodes a nucleotide-binding leucine-rich repeat (NLR) class immune receptor that recognizes its effector, the tobamovirus movement protein (MP). Previously, we found that ToBRFV MP (MP^ToBRFV^) enabled the virus to overcome *Tm-2^2^-* mediated resistance. Yet, it was unknown how *Tm-2^2^* remained durable against other tobamoviruses, such as TMV and ToMV, for over 60 years. Here, we show that the presence of a conserved cysteine (C68) in the MP of TMV (MP^TMV^) is both sufficient to trigger *Tm-2^2^* resistance and essential for viral movement. Substitution of MP^ToBRFV^ amino acid H67 with the corresponding amino acid in MP^TMV^ (C68) activated Tm-2^2^-medited resistance. However, replacement of C68 in TMV and ToMV disabled the infectivity of both viruses. Phylogenetic and structural prediction analysis revealed that C68 is conserved among all *Solanaceae*-infecting tobamoviruses except ToBRFV, and localizes to a predicted jelly-roll fold common to various MPs. Cell-to-cell, and subcellular movement analysis showed that C68 is required for the movement of TMV, by regulating the MP interaction with the endoplasmic reticulum and targeting it to plasmodesmata. The dual role of C68 in viral movement and *Tm-2^2^* immune activation could explain how TMV was unable to overcome this resistance for such a long period.

## 1. INTRODUCTION

Plant viruses cause diseases that severely restrict crop production. Some of the most harmful plant viruses are members of the *Tobamovirus* genus such as Tomato mosaic virus (ToMV) and Tobacco mosaic virus (TMV). Viral symptoms include severe plant yellowing and stunting, reductions in yield and fruit quality and these viruses are highly transmissible by mechanical contact (Broadbent, 1976). The tobamovirus genome consists of a single-stranded, sense RNA of about 6.4 kb that encodes two subunits of an RNA-dependent RNA polymerase (RdRp), a movement protein (MP) and a coat protein (CP) (Goelet et al., 1982). Successful infection depends on the MP, which enables the virus to move from cell-to-cell (Navarro et al., 2019). MP binds to the viral RNA, targets it to plasmodesmata, membrane-lined channels interconnecting adjacent cells, and increases the plasmodesmata size exclusion limit in a process termed ‘gating’ (Reagan and Burch-Smith, 2020). These processes depend on various MP attributes, including its specific subcellular targeting, interaction with cellular factors [e.g., endoplasmic reticulum (ER) and microtubules], and the binding of multiple host proteins (Pitzalis et al., 2018).

Dominant resistance (*R*) genes in plants are pivotal factors for the control of plant pathogens in agriculture. Most *R* genes encode nucleotide-binding leucine-rich repeat (NLR) class of immune receptors that confer resistance by recognizing specific pathogen-derived molecules named effectors (Jones and Dangl, 2006; Adachi et al., 2019). Upon effector recognition, NLRs oligomerize into a structure named ‘resistosome’, to form a calcium-permeable plasma membrane channels (Wang et al., 2019; Martin et al., 2020; Bi et al., 2021). The formed calcium influx then induces programmed cell death of the infected plant tissue that limits pathogen infection in a process termed hypersensitive response (HR) (Meier et al., 2019). The tomato *Tm-2^2^* gene encodes an NLR that confers protection against viruses of the *Tobamovirus* genus, including TMV and ToMV. Tm-2^2^ recognizes and associates with the viral MP, the effector, to trigger the resistance response against tobamoviruses (Weber et al., 1993; Chen et al., 2017). This recognition results in the self-association of Tm-2^2^, which activates HR-mediated immunity against the virus (Wang et al., 2020).

Until recently, *Tm-2^2^* was considered one of the most durable *R* genes in agriculture (Hall, 1980). However, a newly emerged tobamovirus named Tomato brown rugose fruit virus (ToBRFV) has overcome all tobamovirus resistance genes including *Tm-2^2^*, and causes substantial damage to the global tomato industry (Luria et al., 2017; Zhang et al., 2022). Recently, we found that the MP of ToBRFV (MP^ToBRFV^) is the viral factor that enables ToBRFV to overcome *Tm-2^2^* resistance (Hak and Spiegelman, 2021). Expression of MP^ToBRFV^ failed to trigger *Tm-2^2^*-mediated HR and substitution of the original MP sequence of the non-resistance-breaking virus ToMV with MP^ToBRFV^ enabled the recombinant virus to overcome *Tm-2^2^* resistance in tomato. Another recent study mapped the six amino acids in MP^ToBRFV^ (H67, N125, K129, A134, I147, and I168) required for ToBRFV to overcome *Tm-2^2^*, pinpointing the precise sites that enable evasion of *Tm-2^2^*-mediated resistance (Yan et al. 2021).

Since the MP is crucial for intercellular movement, we hypothesized that mutations that enable the MP to overcome *Tm-2^2^*-mediated resistance may also affect cell-to-cell movement of the virus. Our recent findings indicated that MP^ToBRFV^ cell-to-cell movement was reduced compared to MP of TMV (MP^TMV^) (Hak and Spiegelman, 2021). The potential trade-off between *Tm-2^2^* immune evasion and viral movement could explain the high durability of *Tm-2^2^* against TMV and ToMV. However, whether such trade-off exists, and the potential mechanism underlying it are currently unknown. Here, we show that the presence of a cysteine residue (C68) conserved among *Solanaceae*-infecting tobamovirus movement proteins is sufficient to activate *Tm-2^2^* resistance. Our results show that the C68 is required for TMV and ToMV movement by directing MP movement on the ER and subsequent localization to plasmodesmata. Interestingly, in MP^ToBRFV^, the corresponding position is occupied by histidine (H67), but intercellular movement is sustained. The lack of tolerance to mutations in C68 can therefore explain how TMV and ToMV were unable to overcome *Tm-2^2^* for over 60 years.

## 2. RESULTS

### 2.1. The presence of C68 is sufficient to trigger the *Tm-2^2^* resistance response against the tobamovirus MP

Transient co-expression of MP^TMV^ with *Tm-2^2^* in *Nicotiana benthamiana* leaves results in the appearance of HR-mediated cell death due to *Tm-2^2^* immune activation (Figure 1a,b; Hak and Spiegelman, 2021). In contrast, co-expression of MP^ToBRFV^ with *Tm-2^2^* results in only minor cell death (Figure 1a,b; Figure S1). To determine regions of the MP^TMV^ that activate Tm-2^2^, two hybrid MP^TMV^/MP^ToBRFV^ clones were generated. Fusion of MP^TMV^ amino acids 1 to 50 to MP^ToBRFV^ amino acids 51 to 266 (MP^ToBRFV 1-50^/MP^TMV 217-266^) failed to activate Tm-2^2^ (Figure S1). However, MP^TMV^ amino acids 1 to 110 fused to MP^ToBRFV^ amino acids 110 to 266 (MP^TMV 1-110^/MP^ToBRFV 111-266^) triggered Tm-2^2^ response (Figure S1). These results suggested that MP^TMV^ amino acids 51-110 are sufficient to activate Tm-2^2^. Among the MP^ToBRFV^ amino acids required for overcoming *Tm-2^2^*-mediated resistance (Yan et al., 2021), histidine 67 (H67) is the only one that localizes to this region. To test if H67 and the corresponding amino acid in MP^TMV^, cysteine 68 (C68), are determinants of the Tm-2^2^ immune response, we used site-directed mutagenesis to introduce reciprocal mutations in MP^TMV^ and MP^ToBRFV^ thereby changing both amino acids to their identity in the corresponding virus. Indeed, the replacement of MP^ToBRFV^ H67 with cysteine (MP^ToBRFV H67C^) triggered strong HR cell death when co-expressed with *Tm-2^2^* (Figure 1a,b). When H67 was replaced with another amino acid, serine (MP^ToBRFV H67S^), it failed to trigger *Tm-2^2^*-mediated HR cell death, suggesting a specific role for cysteine in *Tm-2^2^* activation (Figure 1a,b). On the other hand, replacement of MP^TMV^ C68 with histidine (MP^TMV C68H^) still triggered HR when co-expressed with *Tm-2^2^* (Figure 1a,b), indicating that MP^TMV^ elements other than C68 are also capable of activating the *Tm-2^2^* response. These results suggest that MP^TMV^ C68 is sufficient, but not essential, to trigger the *Tm-2^2^* immune response.

**Figure 1.**
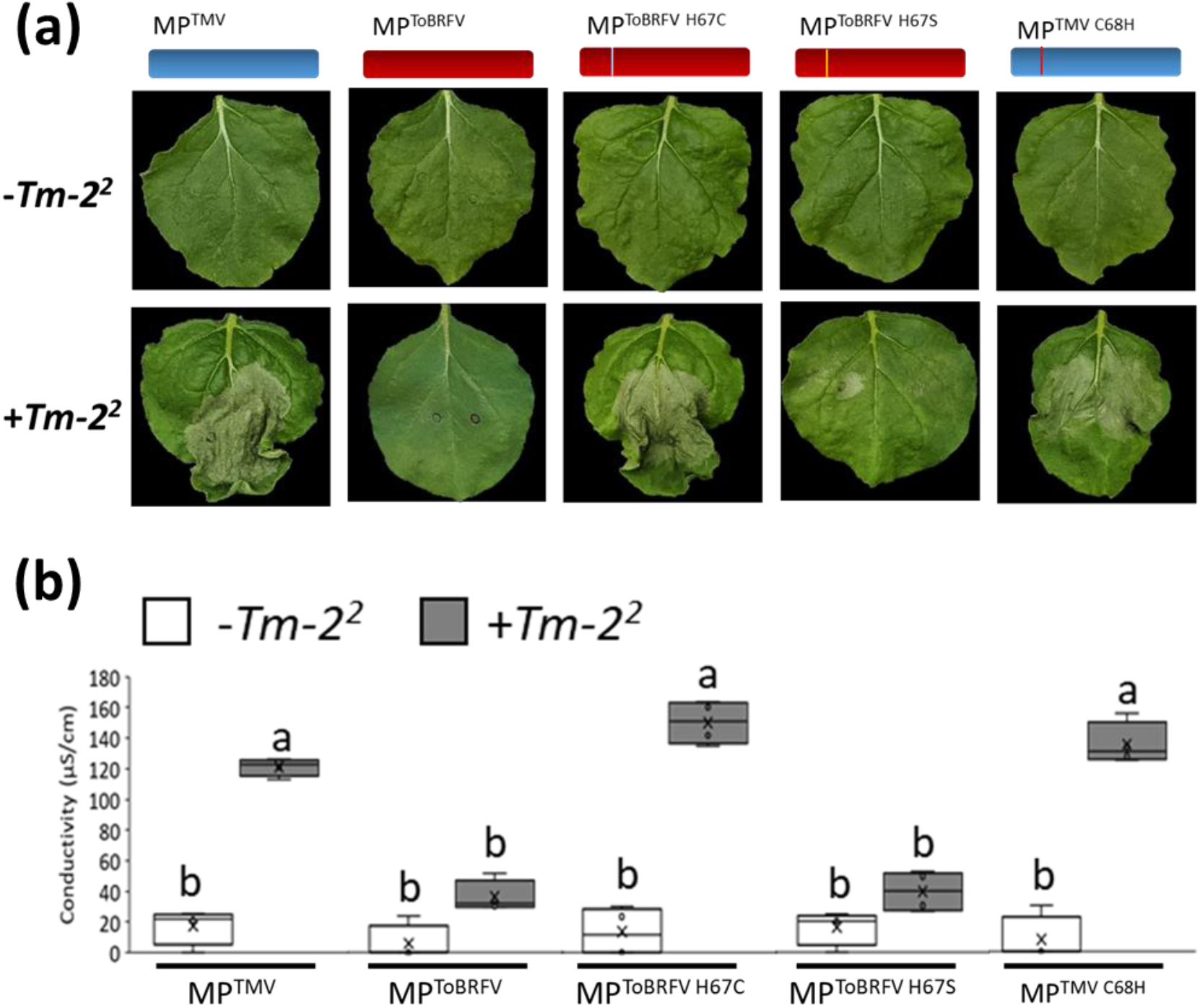
*Tm-2^2^* resistance is activated in the presence of movement protein amino acid C68. (a) Images of *N. benthamiana* leaves transiently expressing MP^TMV^, MP^ToBRFV^, MP^ToBRFV H67C^, MP^ToBRFV H67S^ or MP^TMV C68H^ co-expressed with an empty vector (*-Tm-2^2^*) or with *Tm-2^2^* (+*Tm-2^2^*). Photos were taken 48 hours after infiltration. (b) Electrolyte leakage assay *of N. benthamiana* leaves expressing each construct along with an empty vector (*-Tm-2^2^*) or *Tm-2^2^*(+*Tm-2^2^*). Different letters indicate significant in Tukey’s HSD test (*P*<0.05, *n* ≥ *5*).

### 2.2. MP^TMV^ C68 is conserved in sequence and structure

C68 localizes to a region essential for protein folding (Citovsky et al., 1992) and for association with the ER (Peiro et al., 2014), suggesting its putative function in cell-to-cell movement. To determine the conservation of C68, protein sequence alignment of *Solanaceae*-infecting tobamoviruses was performed (Figure 2a). This analysis revealed that C68 is highly conserved, while H67 is unique to MP^ToBRFV^, suggesting a role for C68 among *Solanaceae*-infecting tobamoviruses.

**Figure 2.**
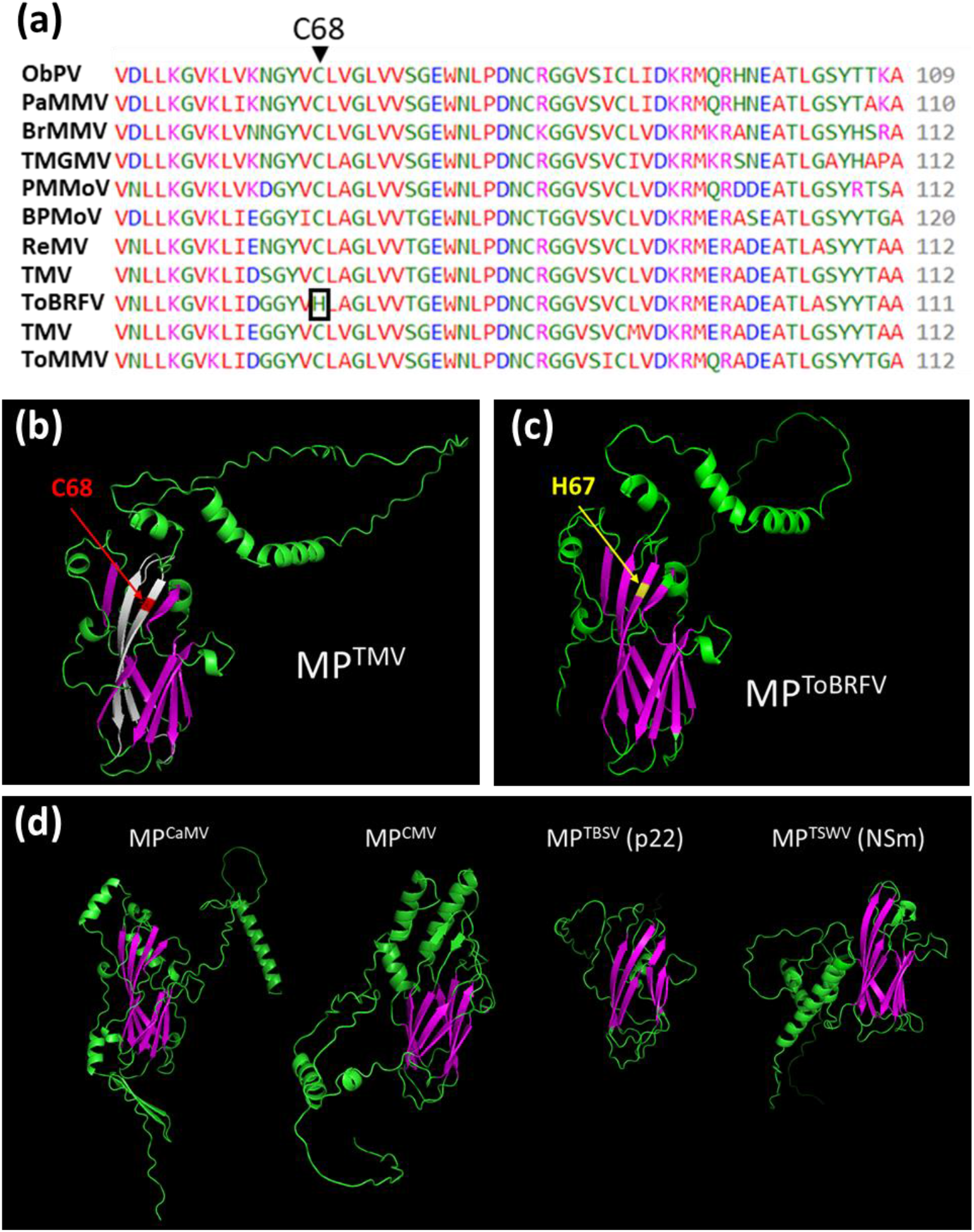
Conservation of the C68 in sequence and structure. **(a)** Protein sequence alignment of solanaceae-infecting tobamovirus movement proteins. C68 is highlighted by black arrow and ToBRFV H67 is highlighted by a black square. **(b-d)** Alphafold models of MP^TMV^ (b), MP^ToBRFV^ (c) and four distant members of the 30K movement protein superfamily (MP^CaMV^, MP^CMV^, MP^TBSV^ and MP^TSWV^) (d). Purple marks the conserved β-barrel fold. White marks in (b) are hydrophobic ER association regions previously described by Peiro et al. (2014). Red arrow marks C68 in MP^TMV^ and yellow marks H67 in MP^ToBRFV^.

To determine the importance of H67/C68, we performed 3D protein structural analysis of MP^TMV^ and MP^ToBRFV^. As there are no existing 3D-structures for any tobamovirus MPs, we used the Alphafold2 modeling platform (Jumper et al., 2021). These analysis provided high-confidence predictions for both proteins, including at the H67/C68 position (Figure 2b,c; Figure S2). Based on this analysis, C68/H67 localized to a jelly roll fold, typical to various viral proteins (Krupovic et al., 2022), which contains two previously-identified membrane interactions motifs (Figure 2b; Peiro et al., 2014). To explore the conservation of this fold, the structure of four other members of the 30K MP superfamily was modeled using Alphafold2: MP of cauliflower mosaic virus (*MP^CaMV^*), MP of cucumber mosaic virus (*MPC^MV^*), MP of tomato bushy stunt virus (MP^TBSV^) and MP of tomato spotted wilt virus (MP^TSWV^) (Figure 2d). Interestingly, this analysis revealed a putative jelly roll fold conserved among all these analyzed MPs. These results suggest that C68 is localized in a MP region conserved in sequence, structure and function, and possibly plays a dual role both in activating the *Tm-2^2^* immune response and cell-to-cell movement.

### 2.3 C68 is essential for TMV and ToMV infection

The conservation of C68 suggests a role for this amino acid in viral infection. To test the function of C68, the C68H mutation was introduced into a ToMV infectious clone (ToMV^MP(C68H)-ToMV^) (Hamamoto et al., 1993). Tomato plants homozygous for the ToMV-sensitive *tm-2* allele or the *Tm-2^2^* resistance allele were then inoculated with the mutant viral RNA (Figure 3a). As controls, plants were also inoculated with non-mutant ToMV and a hybrid version of ToMV harboring MP^ToBRFV^ (ToMV^MP-ToBRFV^) replacing its original MP (Hak and Spiegelman, 2021). Four plants from each treatment were tested three weeks after inoculation for appearance of visual symptoms and presence of viral proteins by immunoblot analysis. As expected, ToMV successfully infected *tm-2* plants, but not *Tm-2^2^* plants (Figure 3a,b). Consistent with our previous report (Hak and Spiegelman, 2021), the ToMV^MP-ToBRFV^ also infected both *tm-2* and *Tm-2^2^* plants (Figure 3a,b). However, ToMV^MP(C68H)-ToMV^ could not infect both genotypes (Figure 3a,b).

**Figure 3.**
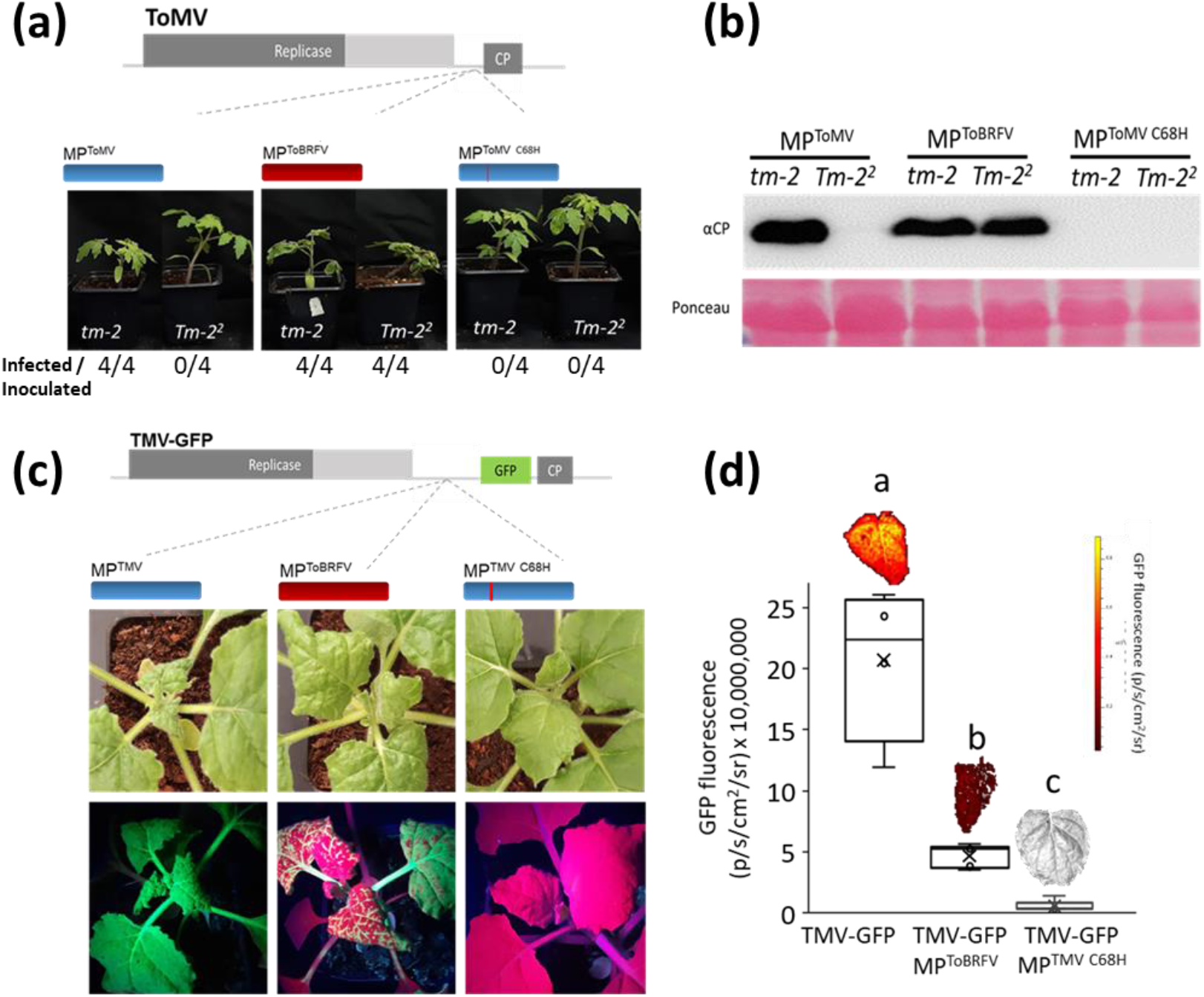
Substitution of MP amino acid C68 results in loss of ToMV and TMV infectivity in tomato and *N. benthamiana*. (a) Tomato plants (cv. Moneymaker) homozygous to the *tm-2* or *Tm-2^2^* allele infected with ToMV, ToMV whose original movement protein was replaced by MP^ToBRFV^ and ToMV with the C68H mutation in the MP ORF. (b) Western-blot analysis of ToMV CP in young leaves of *tm-2* and *Tm-2^2^* plants infected with ToMV expressing the various MPs. Images and leaf samples were taken at 13 days post inoculation (dpi). (c) White light (top) and UV (bottom) images of *N. benthamiana* plants infected with TMV-GFP infectious clones harboring MP^TMV^ (left pane), MP^ToBRFV^ (middle pane) or MP^TMV C68H^ (right pane). (d) *In vivo* imaging-based quantitative analysis of GFP fluorescence in systemic leaves (4^th^ leaf from the apex) of plants infected with TMV-GFP harboring the different movement proteins (MP^TMV^, MP^ToBRFV^ or MP^TMV C68H^). Images were taken at 10 dpi. Different letters indicate significant in Tukey’s HSD test (*P<0.05, n ≥ 4*).

To explore whether the function of C68 is conserved across tobamoviruses and host species, we generated TMV-GFP recombinant virus clones (Lindbo, 2007) containing the MP^ToBRFV^ (TMV-GFP^MP-ToBRFV^) or MP (C68H) mutant (TMV-GFP^MP(C68H)-TMV^) in place of the original TMV MP (Figure 3c). GFP fluorescence levels were monitored in systemic leaves (4^th^ leaf from the apex) using the *in vivo* imaging system (IVIS) (Figure 3d). TMV-GFP and TMV-GFP^MP-ToBRFV^ systemically infected *N. benthamiana* plants (Figure 3C and D). However, TMV-GFP^MP(C68H)-TMV^ was unable to infect *N. benthamiana* plants (Figure 3c,d). These results establish that the amino acid C68 in MP is essential for the systemic infection of both TMV and ToMV in *N. benthamiana* and tomato, respectively. In addition to C68H, we generated other mutations in MP^TMV^ that match their respective identity in MP^ToBRFV^ (Figure S3). These include deletion of valine 5 (ΔV5) and replacement of asparagine 169 with isoleucine (N169I), which is also required for overcoming *Tm-2^2^* (Yan et al., 2021). Neither mutation disabled TMV-GFP from infecting *N. benthamiana* plants (Figure S3), suggesting a unique role for C68 in TMV cell-to-cell movement and therefore infectivity.

### 2.4 Intercellular movement of MP^TMV^ relies on C68

The loss of systemic infection in TMV and ToMV C68H mutants (Figure 3) suggests that C68 is essential for MP function. To determine the effect of C68H on MP cell-to-cell movement, we examined the TMV-GFP inoculation sites in *N. benthamiana* leaves five days after infection (Figure 4). At this time point, the virus is already replicating and moving from cell to cell, resulting in round infection foci. TMV-GFP infection foci size was 1.2 mm^2^ (Figure 4a,d). Consistent with its reduced mobility, TMV-GFP^MP-ToBRFV^ infection foci size was 0. 5 mm^2^ (Figure 4b,d). Strikingly, TMV-GFP^MP(C68H)-TMV^ was able to replicate in the infected cells, but could not move out proximal cells (Figure 4c,d), suggesting that C68 is required for TMV cell-to-cell movement.

**Figure 4.**
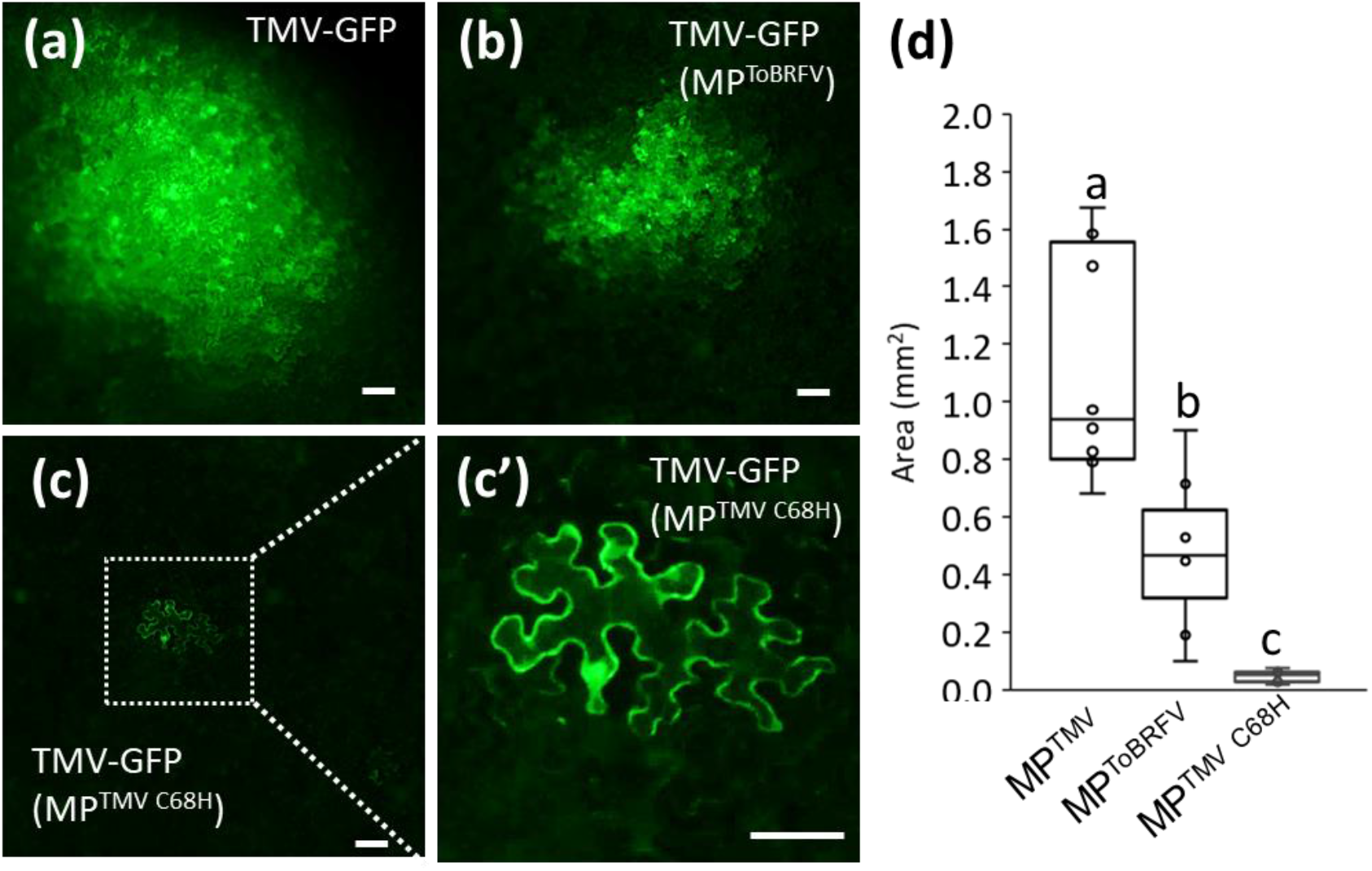
Substitution of MP^TMV^ amino acid C68 results in loss of viral cell-to-cell movement. (a-c) Cell-to-cell movement of the TMV-GFP infectious clones harboring MP^TMV^ (a), MP^ToBRFV^ (b) or MP^TMV C68H^ (c) at 5 dpi. Magnification of the segmented section in the upper panel showing the failure of TMV-GFP MP^TMV C68H^ to move out of the infected cell is shown in (c’). (d) Quantification of infection foci area. Bottom right panel: different letters indicate significant in Tukey’s HSD test (*P*<0.05, *n* ≥ *9*). Scale: 100 μm.

To further examine if C68 is required for MP movement, *MP* ORFs were fused to a gene encoding the yellow fluorescent protein (YFP) and expressed under control of the *35S* promoter. ER-targeted mCherry (ER-mCherry) was expressed from the same binary plasmid as a non-mobile control marking the original cell of expression. *N. benthamiana* leaves were infiltrated with a low dilution of Agrobacterium to express the constructs in distinct epidermal cells as previously described (Hak and Spiegelman, 2021). We observed intercellular movement of both MP^TMV^ and MP^ToBRFV^ (Figure 5a-f,j). However, the C68H mutation abolished cell-to-cell movement of MP^TMV^ (Figure 5g-j). Replacement of other MP^TMV^ amino acids with the corresponding MP^ToBRFV^ amino acids required for overcoming *Tm-2^2^*, N135A and N169I, did not disable intercellular movement of MP^TMV^ (Figure S4), suggesting that C68 plays an important role in controlling the movement function of MP^TMV^.

**Figure 5.**
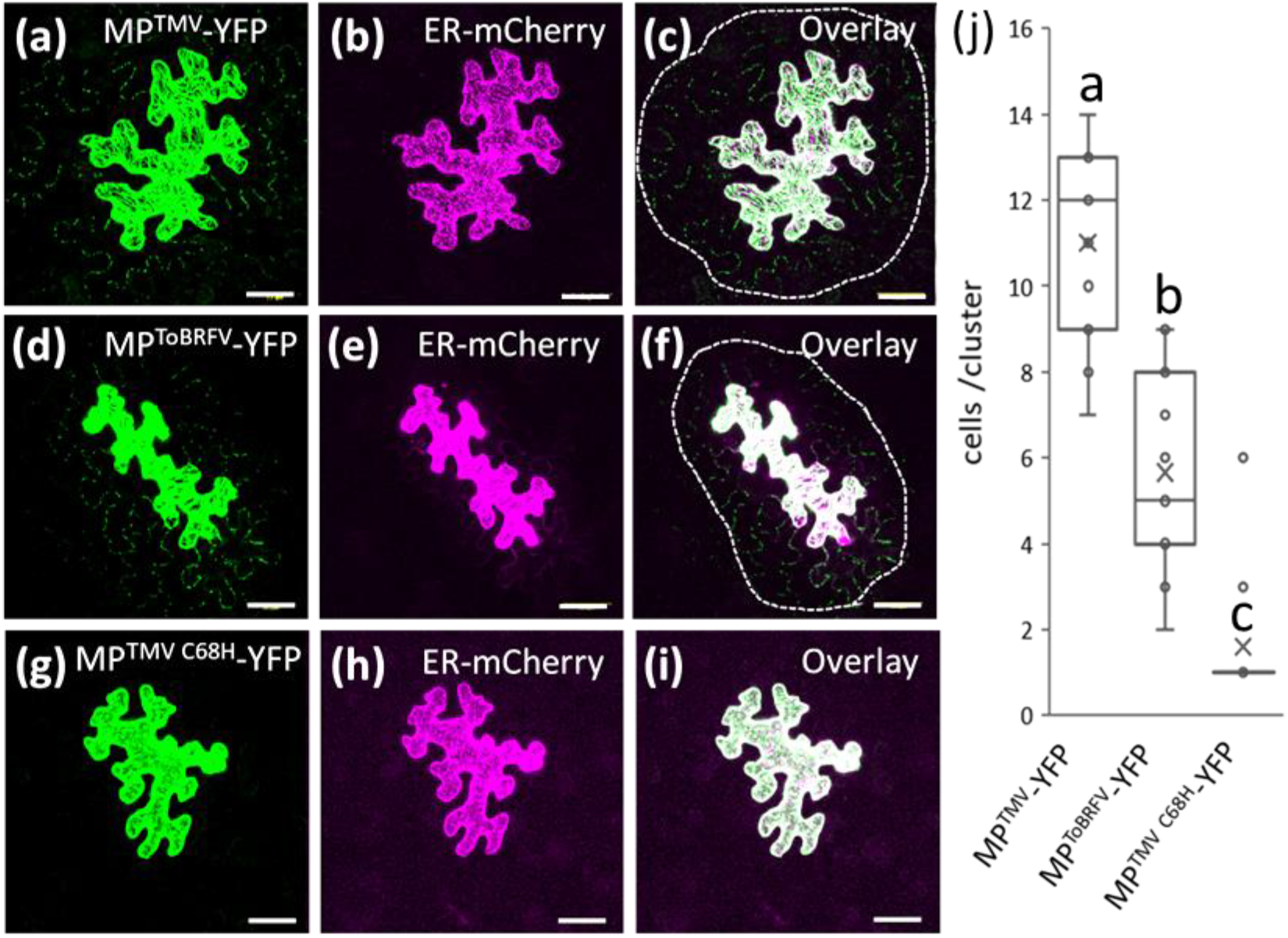
C68 is essential for MP^TMV^ intercellular movement. (a-c) Z-stack maximal projection confocal image of a *N. benthamiana* leaf epidermal cell expressing MP^TMV^-YFP (a), the nonmobile endoplasmic reticulum (ER)-mCherry marker (b) and their overlay (c). (d-f) Z-stack maximal projection confocal image of epidermal cells expressing MP^ToBRFV^-YFP (d), ER-mCherry (e) and their overlay (f). (g-i) Z-stack maximal projection confocal image of epidermal cells expressing MP^TMV^ C^68H^-YFP (g), ER-mCherry (h) and their overlay (i). (j) Quantitative analysis of the number of cells per MP^TMV^-YFP or MP^ToBRFV^-YFP and MP^TMV^ ^C68H^-YFP cluster. Different letters indicate significant in Tukey’s HSD test (P<0.05, *n ≥ 11*). Scale: 50 μm.

### 2.5 C68 determines the localization of MP^TMV^ to the ER and plasmodesmata

To examine how replacement of C68 resulted in the loss of MP^TMV^ cell-to-cell movement, we analyzed the subcellular localization of MP^TMV^ C68H to plasmodesmata using aniline blue callose staining (Figure 6). Specific plasmodesmal localization was calculated as the ratio between MP-YFP fluorescent signal intensity at plasmodesmata (PD) and signal intensity along the plasma membrane (PM). MP^TMV^-YFP showed the highest PD localization (Figure 6a,e). In MP^ToBRFV^-YFP, plasmodesmatal localization was also evident, however, 25% reduced as compared to MP^TMV^ (Figure 6b,e), a result consistent with its reduced cell-to-cell movement. Strikingly, the MP^TMV(C68H)^-YFP mutant was not distinctly targeted to plasmodesmata (Figure 6c,e). Instead, this protein localized to the plasma membrane, as observed by co-localization with the Flot1-RFP plasma membrane marker (Figure 6d,e). These result indicate that C68 is required for the MP^TMV^ intercellular transport function by targeting it to plasmodesmata.

**Figure 6.**
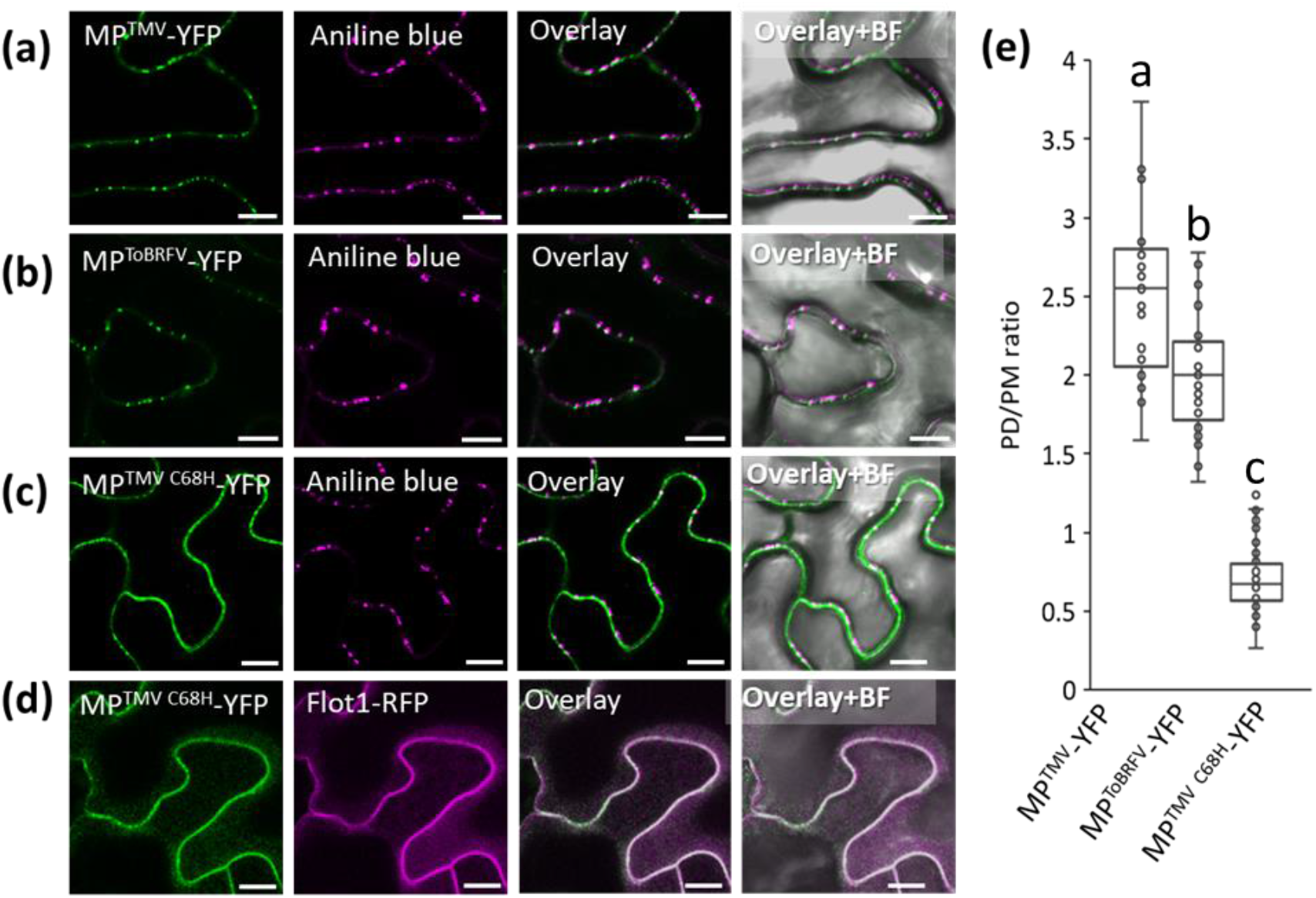
C68 is essential MP^TMV^ plasmodesmal localization. Localization of MP^TMV^-YFP (a), MP^ToBRFV^-YFP (b) and MP^TMV C68H^-YFP (c) proteins (green) To plasmodesmata (purple). Plasmodesmata were labeled using aniline blue callose staining. Note that the MP^TMV C68H^-YFP (c) fails to specifically be targeted to plasmodesmata. (C) Quantitative analysis of the accumulation of the various MP-YFP proteins in plasmodesmata. Calculation was based on the ratio between MP-YFP signal in plasmodesmata (PD) and the average fluorescent signal in the plasma membrane (PM). Different letters indicate significant in Tukey’s HSD test (*P<0.05, n ≥ 25*). Scale: 10 μm.

As shown in Figure 2, C68 is part of an ER interaction motif. Since targeting of MP to plasmodesmata requires its interaction with the ER network (Wright et al., 2007), we postulated that the C68H mutation may alter MP^TMV^ interaction with the ER. To address this, we used confocal microscopy to image the upper plane of *N. benthamiana* epidermal cells co expressing ER-mCherry with either MP^TMV^-YFP or MP^TMV(C68H)^-YFP (Figure 7). In these regions, MP^TMV^-YFP formed vesicles that were 76% co-localized with the ER (Figure 7a-c,g). On the other hand, the localization of MP^TMV(C68H)^-YFP was altered to microdomain-like structures, and the co-localization with the ER was calculated at only 48% (Figure 7b-g). We further used live confocal imaging assessed the ability of the MP to traffic on the ER network. The MP^TMV^-YFP-associated vesicles were largely mobile, with an average velocity of 1.48 μm/sec (Figure 7h-i; Video S1). In marked contrast, MP^TMV(C68H)^-YFP bodies were largely immobile (Figure 7h-i; Video S2). These results suggest that C68 is required for MP^TMV^ targeting to plasmodesmata by regulating its association with the ER.

**Figure 7.**
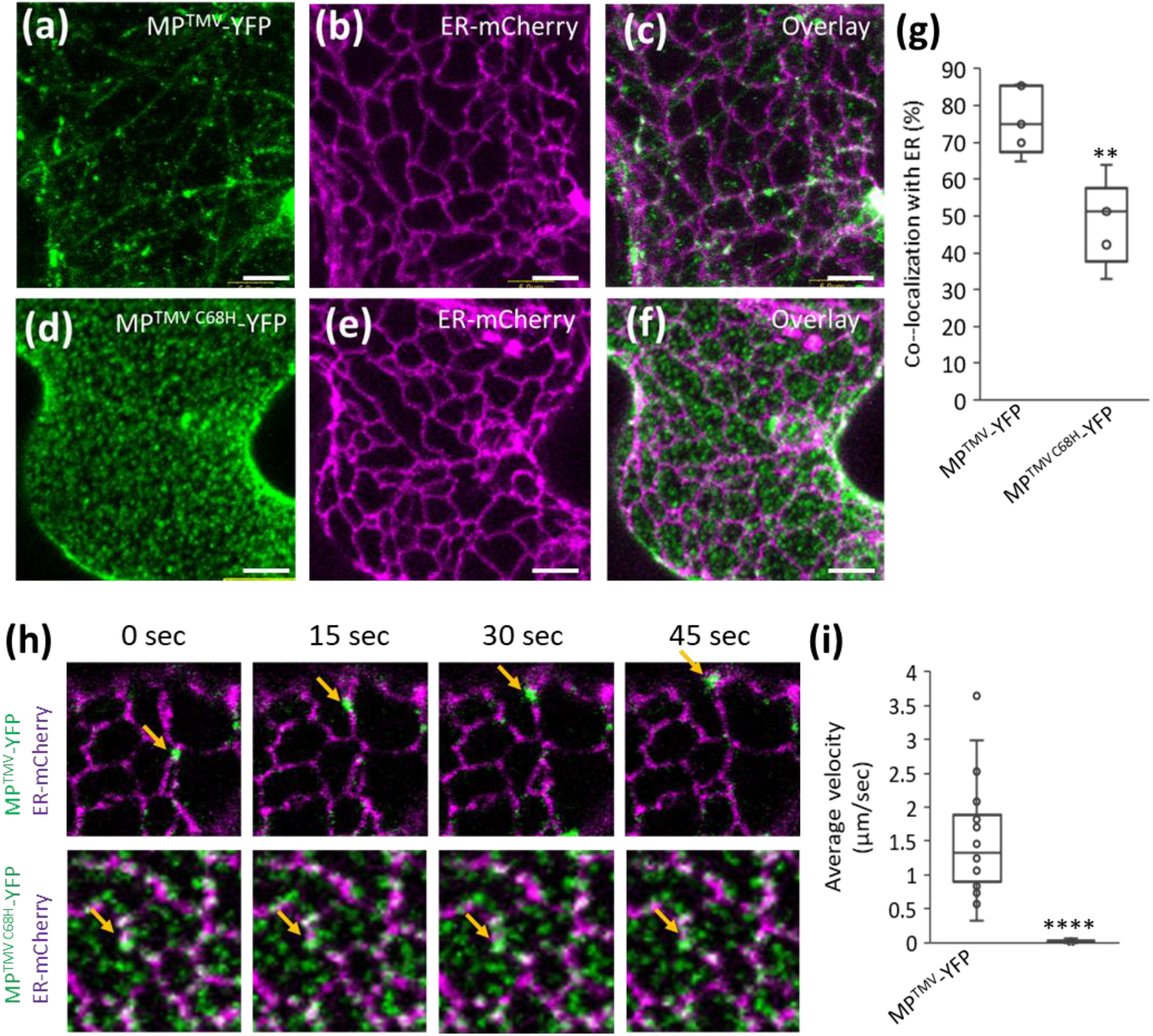
C68 is required for MP^TMV^ interaction with the ER and intracellular movement. Surface imaging of upper *N. benthamiana* leaf epidermal cell planes. (a-c) Expression of MP^TMV^-YFP (a), ER-mCherry (b) and an overlay of both proteins (c). (d-f) expression of MP^TMV C68H^-YFP (d), ER-mCherry (e) and an overlay of both proteins (f). (g) Quantitative analysis of MP^TMV^-YFP and MP^TMV C68H^-YFP co-localization with ER-mCherry. (h) Dynamics of MP^TMV^-YFP (top) and MP^TMV C68H^-YFP (bottom) on the ER network. Note that while MP^TMV^-YFP are mobile on the ER, MP^TMV C68H^-YFP are immobile (yellow arrow). (i) Quantification of MP^TMV^-YFP and MP^TMV C68H^-YFP vesicle velocity. Scale = 5 μm. (** *p<0.01, **** p<0.0001* in student’s t-test, *n ≤ 24*).

## 3. DISCUSSION

The co-evolution of *R* genes and their effectors (also known as *Avr* factors) is a major driving force in the arms race between plants and pathogens. Durability of *R* genes relies on the importance and conservation of its targeted effector (Jones and Dangl, 2006). A successful *R* gene will therefore target effector elements essential for pathogen infectivity and/or survival. However, there are only a few known examples for the mechanisms underlying such trade-offs. Previously, we have shown that overcoming *Tm-2^2^* by MP^ToBRFV^ was associated with an attenuation of viral movement (Hak and Spiegelman, 2021), suggesting that the elements required for evading *Tm-2^2^* also function in MP intercellular movement. Here we show that among the amino acids required to overcome *Tm-2^2^* (Yan et al., 2021), H67 localizes to a critical site for MP function, replacing a cysteine (C68) conserved among *Solanaceae*-infecting tobamoviruses, and placed within a structure conserved among multiple movement proteins. Substitution of MP^ToBRFV^ H67 with cysteine triggered *Tm-2^2^* resistance, suggesting that the presence of this amino acid is sufficient to activate Tm-2^2^. This is consistent with a previous finding that H67C substitution disabled the capacity of ToBRFV to systemically infect plants harboring *Tm-2^2^* (Yan et al., 2021). Interestingly, our results show that the same cysteine is essential for the function of ToMV and TMV, by enabling MP subcellular targeting to plasmodesmata via the ER.

The ER plays a major role in TMV intercellular transport (Pitzalis et al., 2018). Initially, TMV replication complexes are associated with the ER. Then, MP^TMV^-associated complexes detach and move along the ER membrane to be targeted to plasmodesmata (Wright et al., 2007; Sambade et al., 2008; Guenoune-Gelbart et al., 2008). The interaction of MP^TMV^ is mediated by two hydrophobic regions, which associate, but do not integrate into the ER membrane. These regions span between amino acids 61-80 and 148-167. Although sequentially distant, our Alphafold-based homology model predicted these two sections to be topologically interacting, and take part in a putative jelly-roll fold common to various MPs (Figure 2b-d). These results suggest that Tm-2^2^ targets the viral MP in an element crucial for its function.

There are only few examples in which resistance-breaking mutations cause penalties in viral infectivity or competition (García-Arenal and Fraile, 2013). Mutations in the Potato virus Y Nla protein that enable it to overcome the *Ry*-mediated resistance also caused loss of its protease activity (Mestre et al., 2003). Mutations in Turnip mosaic virus which overcome TuRB01 and TuRB04 resistances in *Brassica napus* impaired the virus’s ability to compete with a non-resistance-breaking viral strain on a susceptible host (Jenner et al., 2002). *Tm-1* resistance-breaking mutations in the ToMV replicase delay viral replication in tobacco protoplasts (Ishibashi et al., 2012). Mutations in Pepper mild mottle virus CP that break *L*-mediated resistance in pepper were associated with high fitness penalties (Fraile et al., 2011). However, recently it was shown that the effect of resistance-breaking mutations is more complex and depends on the specific mutation, the host genotype, and the infecting isolate (Moreno-Pérez et al., 2016).

Hall (1980) suggested that the durability of *Tm-2^2^*-mediated resistance against TMV and ToMV was a result of the reduced capacity of these viruses to evolve to overcome *Tm-2^2^*, and predicted that this mutation would likely alter fundamental viral characteristics. Our finding that overcoming *Tm-2^2^* requires the substitution of C68, which is essential for viral movement, supports this hypothesis. In light of these findings, how ToBRFV was able to overcome this resistance, whereas TMV and ToMV failed to do so? Previous lineage analysis predicted that ToBRFV likely evolved on an unknown host other than tomato, and invaded tomato due to a host-shifting event (Maayan et al., 2018). This allowed MP^ToBRFV^ to diverge from MP^TMV^ and MP^ToMV^ in multiple elements, including C68/H67, while sustaining viral movement. On the other hand, in TMV and ToMV, loss of C68 would require additional compensatory mutations for MP to retain functionality. The requirement of multiple mutations to overcome *Tm-2^2^* combined with high levels of fitness penalties would be highly unlikely or rare and uncompetitive.

Collectively, we find that the conversion of cysteine at position 67 (H67) in MP^ToBRFV^ is crucial for evading *Tm-2^2^* resistance. However, in MP^TMV^, substitution of cysteine in the corresponding position (C68) results in the abolishment of cell-to-cell movement. We conclude that this strong trade-off between viral movement and breaking host resistance could be one of the primary reasons for the high durability of *Tm-2^2^* against TMV and ToMV for over 60 years.

## 4. EXPERIMENTAL PROCEDURES

### 4.1 Plant materials

Tomato (*Solanum lycopersicum* L.) seeds (cv. Moneymaker) homozygous for the *tm-2* allele (LA2706) or the *Tm-2^2^* allele (LA3310) were obtained from the Tomato Genome Research Center, University of California, Davis. *Nicotiana benthamiana* and tomato plants were grown in soil in a light- and temperature-controlled chamber at 25°C in a 16-h photoperiod. Five-week-old *N. benthamiana* plants were used for agroinfiltration and three-week-old tomato plants were used for mechanical inoculation of ToMV.

### 4.2 Plasmid construction

Plasmids *p35S:Tm-2^2^, p35S:MP^TMV^, p35S:MP^ToBRFV^, p35S:MP^TMV^-YFP* + *ER-mCherry* and *p35S:MP^ToBRFV^-YFP* + *ER-mCherry* were assembled by Golden Gate cloning using the MoClo tool kit for plants (Addgene) (Weber et al., 2011) as described (Hak and Spiegelman, 2021). Generation of the vectors TMV-GFP^MP-ToBRFV^ and ToMV^MP-ToBRFV^, based on the pJL24 (Lindbo, 2007) and pTLW3 (Hamamoto et al., 1993) were described previously (Hak and Spiegelman, 2021). Hybrid movement proteins were generated using fusion PCR. All the constructs that contain point mutations were made using site-directed mutagenesis (Ho et al., 1989) and confirmed by sequencing.

### 4.3 Transient expression and agroinfiltration

*Agrobacterium tumefaciens* strain EHA105 harboring a binary vector was grown overnight at 28°C. Cell cultures were resuspended in 2-morpholinoethane sulfonic acid (MES) buffer (10 mM MgCl_2_, 10 mM MES, 150 μM acetosyringone, pH 5.6) to OD_600_ = 0.5. For MP-YFP and TMV-GFP cell-to-cell movement assays, OD_600_ = 0.0001 and OD_600_ = 5.00E^-6^ were used, respectively. Agrobacterium solutions were infiltrated using a needleless syringe into the abaxial side of the fourth or fifth leaf of *N. benthamiana* plants.

### 4.4 Electrolyte leakage assay

*p35S:MP* constructs were agroinfiltrated into *N. benthamiana* leaves (*n = 4*) at OD_600_ = 0.25 at a 1:1 ratio with either *p35S:Tm-2^2^* or an empty vector. Twenty-seven hours after agroinfiltration, four leaf discs (1 cm diameter) were cut from each leaf and washed for 10 to 30 min in distilled water. The leaf discs were then carefully transferred into a tube containing 4 ml of distilled water and placed on a low speed orbital shaker. Electric conductivity was measured after 6 hours, using a benchtop conductivity meter. Each experiment was repeated at least 3 times.

### 4.5 TMV-GFP infection and quantification

For systemic infection experiments, *A. tumefaciens* harboring TMV-GFP or TMV-GFP^MP-ToBRFV^ vectors at OD_600_ = 0.25 were infiltrated at a 1:1 ratio with the silencing suppressor p19 into the fourth leaf of 4-week-old *N. benthamiana* plants. GFP fluorescence was monitored 6 days post-infiltration. Images were acquired and analyzed using an IVIS Lumina LT (PerkinElmer) equipped with a XFOV-24 lens and Living Image 4.3.1 software (PerkinElmer) set (excitation/emission: 420 nm/520 nm). The optical luminescent image data were displayed in pseudocolor that represents intensity in terms of radiance (photons per second per square centimeter per steradian) and were calculated as average radiance per leaf. For cell-to-cell movement of TMV-GFP, imaging of GFP was obtained and analyzed at 5 dpi using a Nikon SMZ-25 stereomicroscope equipped with a Nikon-D2 camera and NIS Elements v. 5.11 software. The number of plants for each experiment was n=4, and experiments were repeated at least 3 times.

### 4.6 In vitro transcription and infection of ToMV

Two microliters of ToMV pTLW3 plasmid, ToMV^MP-ToBRFV^, or ToMV^MP-TMV^(C^68H^) were linearized with *SmaI* and were prepared by a gel extraction kit (Zymo Research). In vitro transcription was performed using the mMESSAGE mMACHINE T7 kit (Invitrogen by Thermo Fisher Scientific). Two drops of 2.5 μl of transcript were used to mechanically inoculate *N. benthamiana* plants dusted with carborundum powder prior to inoculation. Leaves of infected plants were collected 5 to 7 days post inoculation to serve as inoculum for 3-week-old tomato plants.

### 4.7 Detection of ToMV by immunoblotting

For detection of the systemic spread of the virus in tomato, the first young leaf (10 to 30 mm) and apex were collected and ground, while frozen, in a microcentrifuge tube. Laemmli buffer (40 μl of 3×) (100 mM Tris, 2% SDS, 20% glycerol, 4% β-mercaptoethanol, pH 6.8) was added and mixed into the sample, followed by centrifugation for 10 min and boiling of the supernatant for 5 min. Samples were run on 12% SDS-PAGE acrylamide gels, and transferred to nitrocellulose membranes (Protran), and were blocked with 3% skimmed milk in Tris buffer saline-Tween. CP of the virus was detected by rabbit anti-tobamovirus CP (1:20,000 [courtesy of A. Dombrovsky]), and anti-rabbit horseradish peroxidase (1:20,000 [Jackson Immunoresearch]). Chemiluminescence was observed using Elistar Supernova as substrate (Cyanagen), and images of protein bands were acquired using the Alliance UVITEC software.

### 4.8 Cell-to-cell movement analysis of MP-YFP

Experiments were performed as previously described (Hak and Spiegelman, 2021). Constructs expressing both MP-YFP and ER-mCherry were agroinfiltrated into *N. benthamiana* and visualized 36 h later. For confocal imaging, we used an Olympus IX 81 inverted laser scanning confocal microscope (Fluoview 500) equipped with an OBIS 488 and 561 nm laser lines and a 60× 1.0 NA PlanApo water immersion objective. YFP and mCherry were excited at 488 and 561 nm and were imaged using BA505-525 nm and BA575-620 nm emission filters, respectively. The number of YFP-fluorescent cells surrounding the original cell of expression was quantified to determine the level of movement for each MP.

### 4.9 Subcellular localization of MP-YFP

For ER localization, double expression plasmids harboring MP-YFP and ER-mCherry were used (Hak and Spiegelman, 2021). For plasma membrane localization, MP-YFP were co-expressed with Flotilin1 fused to RFP under the 35s promoter (Flot1-RFP) (Pizarro et al., 2019). For plasmodesmatal localization, aniline blue solution (0.1% in water) was infiltrated into the abaxial side of the leaf and 10 minutes later imaged. Images wer taken using Olympus IX 81 inverted laser scanning confocal microscope (Fluoview 500). Excitation of analinie blue at 405 nm was done using a BA430–460 filter with an UPlanSApo 60× 1.35 oil objective, 8/0.17 FN26.5. Quantification of MP-YFP accumulation intensity in plasmodesmata, plasma membrane and ER was performed using the ImageJ software (https://imagej.nih.gov/ij/) by measuring the YFP signal intensity co-localizaed with each of the markers.

### 4.10 Multiple sequence alignment

Sequence alignment of the *Solanaceae*-infecting tobamoviruses MPs pepper mild mottle virus (NP_619742.1), bell pepper mottle virus (YP_001333652.1), rehmannia mosaic virus (YP_001041891.1), TMV (NP_597748.1), ToBRFV (YP_009182170.1), ToMV (NP_078448.1), tomato mottle mosaic virus (YP_008492930.1), obuda pepper virus (NP_620843.1), paprika mild mottle virus (NP_671720.1), brugmansia mild mottle virus (YP_001974325.1) and tobacco mild green mosaic virus (sp|P18338|) was performed using Clustal Omega (https://www.ebi.ac.uk/Tools/msa/clustalo/0).

### 4.11 Protein structure prediction using Alphafold2

The 3D-structure of MP^TMV^ (NP_597748.1), MP^ToBRFV^ (YP_009182170.1) MP^CaMV^ (NP_056724.1), MP^CMV^ (NP_040776.1), MP^TBSV^ (NP_062900.1) and MP^TSWV^ (AEK06236.1) were predicted using the artificial Intelligence-based Alpha-Fold2 through the AlphaFold Colab notebook platform (Jumper et al., 2021). For the structural comparisons, we used only the predicted model with the highest confidence score computed by AlphFold for each protein. PyMOL Molecular Graphics System (Version 2.4, Schrödinger) was used to examine the structural models and generate images.

## SUPPLEMENTAL DATA

**Figure S1.** MP^TMV^ amino acids 51-110 activate *Tm-2^2^* resistance.

**Figure S2.** High confidence for Alphafold modeling of MP^TMV^ and MP^ToBRFV^.

**Figure S3.** Mutagenesis of MP^TMV^ amino acids other than C68 to match their respective identity in MP^ToBRFV^ does not result in loss of TMV infectivity.

**Figure S4.** Cell-to-cell movement of MP^TMV^-YFP with substitution mutations to corresponding MP^ToBRFV^ amino acids required to overcome *Tm-2^2^*.

**Video S1.** Trafficking of MP^TMV^-YFP vesicles on the ER

**Video S2.** Disabled trafficking of MP^TMV C68H^-YFP punctae on the ER

## ACKNOWLEDGEMENTS

We thank M. Heinlein (French National Centre for Scientific Research), B. Falk (University of California Davis) and M. Ishikawa (Japanese National Agriculture and Food Research Organization) for providing the for providing the TMV-GFP and ToMV infectious clones. We thank A. Dombrovsky for anti-tobamovirus CP antibody and I. Levin (Argiculture Research Organization, Israel) for the tomato (cv. Moneymaker) lines used in the manuscript. We thank V. Gaba (Argiculture Research Organization, Israel) for critical reading of this manuscript.

## AUTHOR CONTRIBUTIONS

H.H, S.P.D.K and Z.S conceptualized and designed the different experiments. H.H cloned the different constructs, and performed all virus and cell-to-cell transport assays described in the manuscript. The hypersensitive response assays were performed by H.H and Y.S. H.R S.S. conceived and performed the protein modeling using Alphafold2. Z.S wrote this manuscript in collaboration with S.P.D.K.

## DATA AVAILABILITY

The data that support the findings of this study are available from the corresponding author upon reasonable request.

