## Supplementary material for "*Tm-2^2^* resistance targets a conserved cysteine essential for tobacco mosaic virus (TMV) movement": Figures S1-S4

### Figure S1

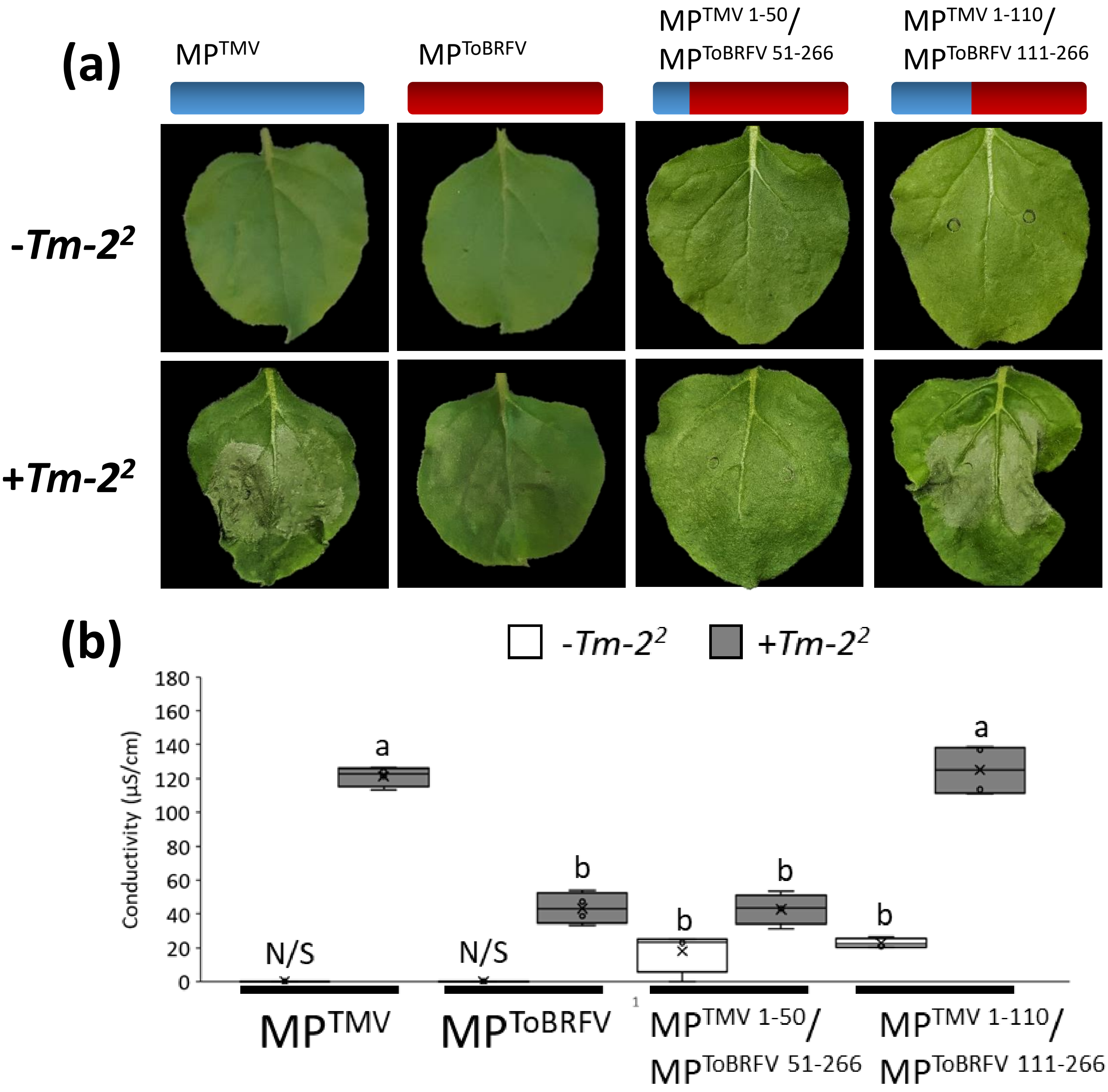


**Figure S1. MP$^{TMV}$ amino acids 51-110 activate *Tm-2*$^2$ resistance.** (a) Cell death responses in *N. benthamiana* leaves expressing MP$^{TMV}$, MP$^{ToBRFV}$, and the hybrid MPs MP$^{TMV\ 1\text{-}50}$/ MP$^{ToBRFV\ 51\text{-}266}$ and MP$^{TMV\ 1\text{-}110}$/ MP$^{ToBRFV\ 111\text{-}266}$. The various MPs co-expressed with an empty vector (-*Tm-2*$^2$) or with *Tm-2*$^2$ (+*Tm-2*$^2$). (b) Electrolyte leakage assay *of* MP$^{TMV}$, MP$^{ToBRFV}$ and the hybrid MPs MP$^{TMV\ 1\text{-}50}$/ MP$^{ToBRFV\ 51\text{-}266}$ and MP$^{TMV\ 1\text{-}110}$/ MP$^{ToBRFV\ 111\text{-}266}$ expressed with an empty vector (-*Tm-2*$^2$) or with *Tm-2*$^2$ (+*Tm-2*$^2$). Different letters indicate significant in Tukey's HSD test ($P<0.05$, $n \geq 4$).

Figure S2

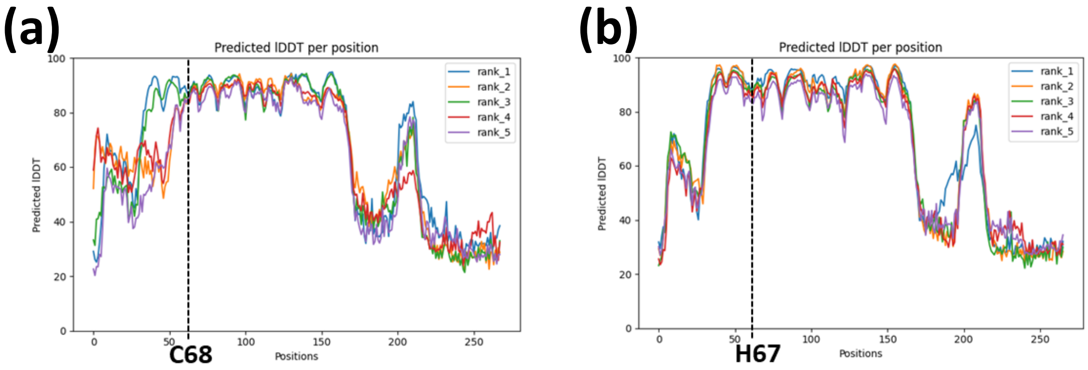


**Figure S2. High confidence for Alphafold modeling of $MP^{TMV}$ and $MP^{ToBRFV}$.** Predicted local distance difference test for $MP^{TMV}$ (a) and $MP^{ToBRFV}$ indicating high level of structure confidence for amino acids 40-171, including C68 and H67, respectively (broken line).

Figure S3

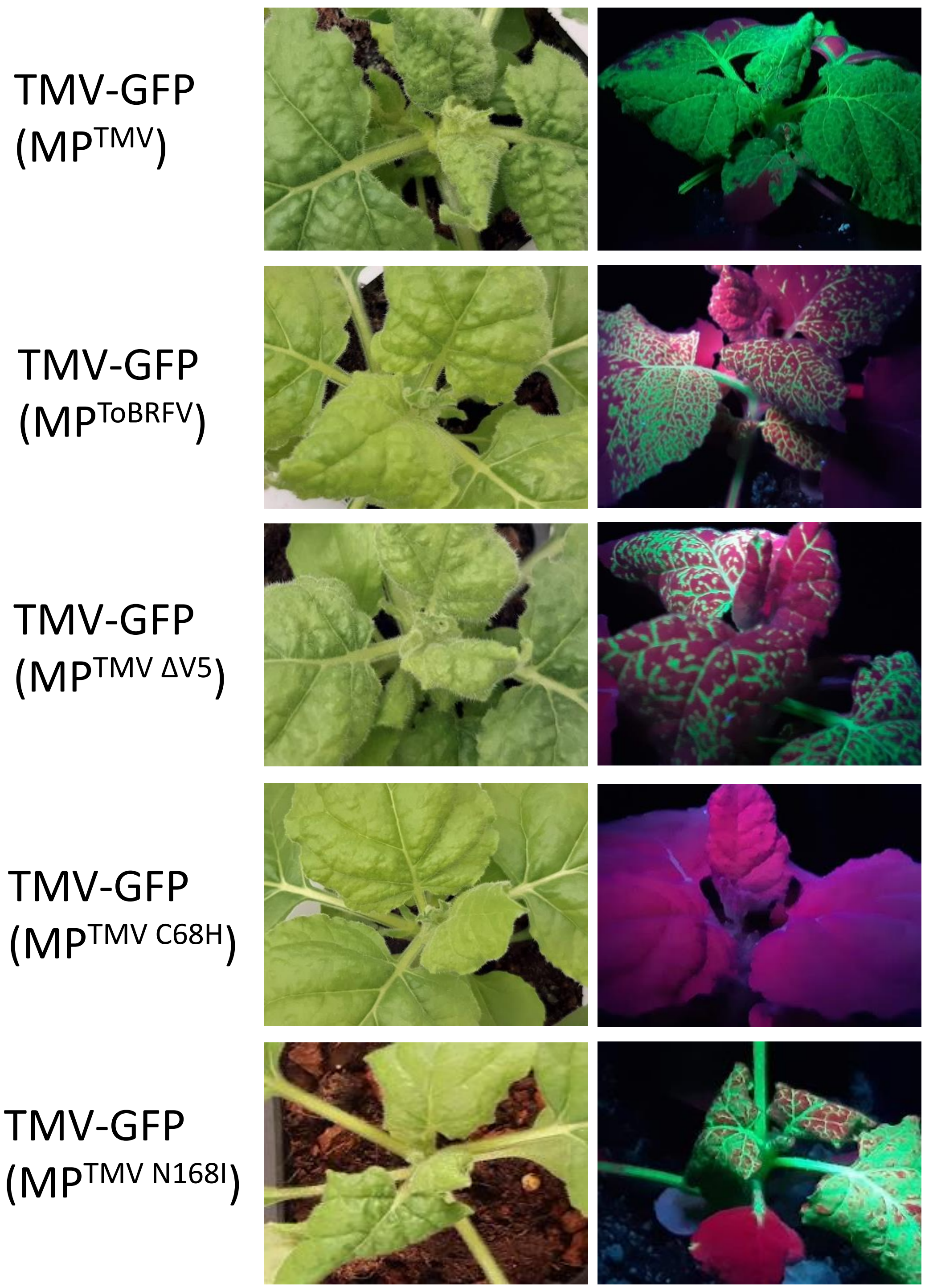


**Figure S3. Mutagenesis of MP$^{TMV}$ amino acids other than C68 to match their respective identity in MP$^{ToBRFV}$ does not result in loss of TMV infectivity.** White light (left) and UV (right) images of *N. benthamiana* plants infected with TMV-GFP infectious clones harboring MP$^{TMV}$, MP$^{ToBRFV}$ or MP$^{TMV}$ with mutations to their respective identity in MP$^{ToBRFV}$: ΔV5, C68H and N169I. The only mutation disabling TMV infectivity was C68H.

### Figure S4

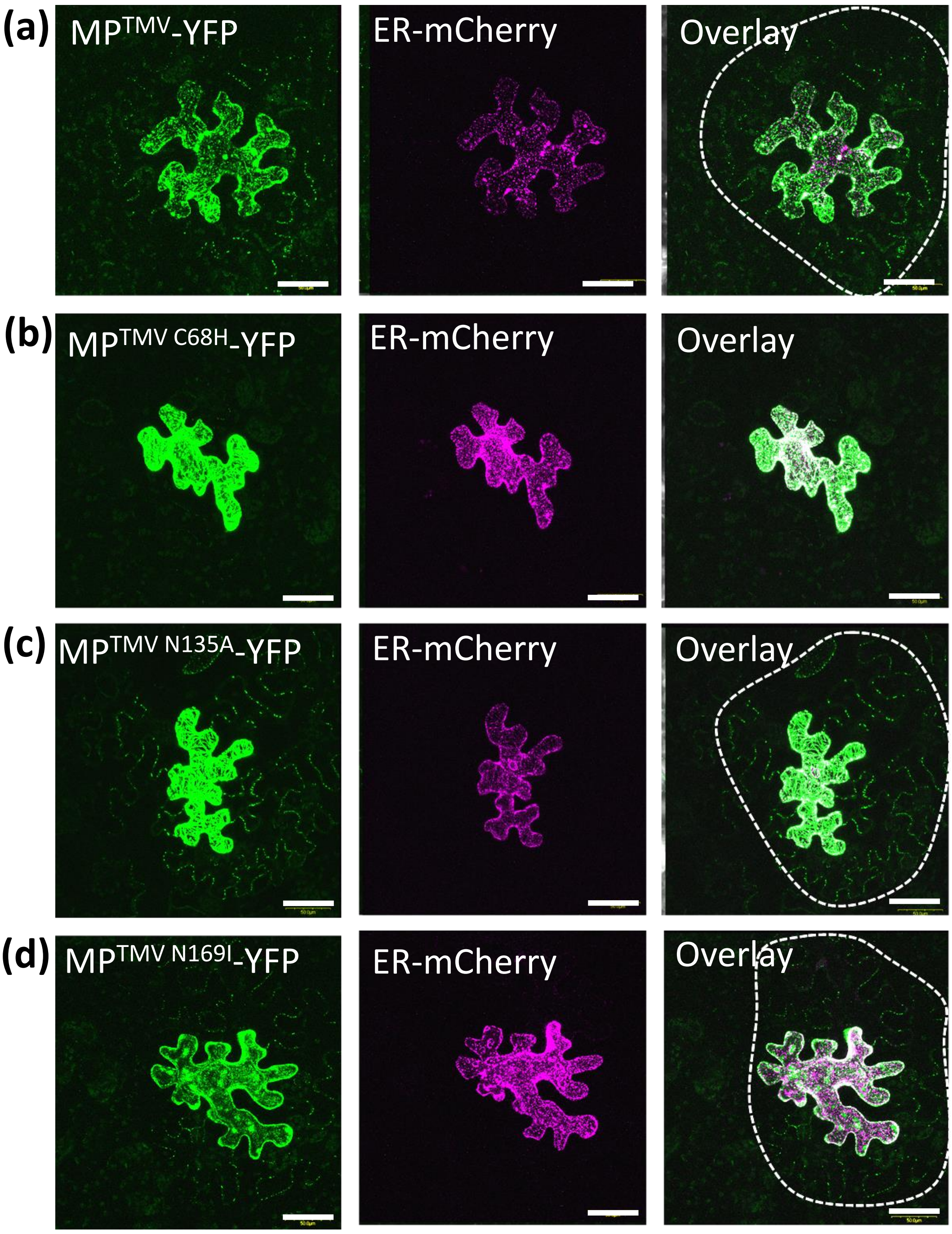


**Figure S4. Cell-to-cell movement of $MP^{TMV}$-YFP with substitution mutations to corresponding $MP^{ToBRFV}$ amino acids required to overcome *Tm-2²*.** Z-stack maximal projection confocal image of *Nicotiana benthamiana* leaf epidermal cells expressing $MP^{TMV}$-YFP (left panel) with no mutations (a) and mutations to corresponding $MP^{ToBRFV}$ amino acids required for overcoming Tm-22: C68H (b), N135A (c) and N169I (Yan et al., 2021). The nonmobile endoplasmic reticulum (ER)-mCherry served as negative control for protein movement (middle panel). Overlay of $MP^{TMV}$-YFP and ER-mCherry signals is presented at the right panel. Note that the only mutation that restricted cell to cell movement is C68H. Scale = 50 µm
